# Large-scale sequence similarity analysis reveals the scope of sequence and function divergence in PilZ domain proteins

**DOI:** 10.1101/2020.02.11.943704

**Authors:** Qing Wei Cheang, Shuo Sheng, Linghui Xu, Zhao-Xun Liang

## Abstract

PilZ domain-containing proteins constitute a superfamily of widely distributed bacterial signalling proteins. Although studies have established the canonical PilZ domain as an adaptor protein domain evolved to specifically bind the second messenger c-di-GMP, mounting evidence suggest that the PilZ domain has undergone enormous divergent evolution to generate a superfamily of proteins that are characterized by a wide range of c-di-GMP-binding affinity, binding partners and cellular functions. The divergent evolution has even generated families of non-canonical PilZ domains that completely lack c-di-GMP binding ability. In this study, we performed a large-scale sequence analysis on more than 28,000 single- and di-domain PilZ proteins using the sequence similarity networking tool created originally to analyse functionally diverse enzyme superfamilies. The sequence similarity networks (SSN) generated by the analysis feature a large number of putative isofunctional protein clusters, and thus, provide an unprecedented panoramic view of the sequence-function relationship and function diversification in PilZ proteins. Some of the protein clusters in the networks are considered as unexplored clusters that contain proteins with completely unknown biological function; whereas others contain one, two or a few functionally known proteins, and therefore, enabling us to infer the cellular function of uncharacterized homologs or orthologs. With the ultimate goal of elucidating the diverse roles played by PilZ proteins in bacterial signal transduction, the work described here will facilitate the annotation of the vast number of PilZ proteins encoded by bacterial genome and help to prioritize functionally unknown PilZ proteins for future studies.

**Importance:** Although PilZ domain is best known as the protein domain evolved specifically for the binding of the second messenger c-di-GMP, divergent evolution has generated a superfamily of PilZ proteins with a diversity of ligand or protein-binding properties and cellular functions. We analysed the sequences of more than 28,000 PilZ proteins using the sequence similarity networking (SSN) tool to yield a global view of the sequence-function relationship and function diversification in PilZ proteins. The results will facilitate the annotation of the vast number of PilZ proteins encoded by bacterial genomes and help us prioritize PilZ proteins for future studies.

## Introduction

C-di-GMP is an intracellular nucleotide messenger found in many environmental and pathogenic bacteria. High cellular c-di-GMP concentration is generally correlated with the sessile lifestyle, production of extracellular polysaccharides (EPS) and formation of surface-associated biofilm. C-di-GMP exerts its biological effect by binding to a wide variety of protein and riboswitch effectors. PilZ domain is the first discovered c-di-GMP binding protein domain and arguably the most prevalent one ^1–3^. The PilZ domain is characterized by a seven or eight-strand β-barrel consisting of two antiparallel β-sheets and a long and flexible N-terminal loop. Canonical PilZ domains bind monomeric or dimeric c-di-GMP with a wide range of affinities using two key motifs: the arginine-rich motif (RxxxR) from the N-terminal loop and the (D/N)xSxxG motif from the β-strands ^4–7^. Apart from the canonical PilZ domains, non-canonical PilZ domain proteins that lack c-di-GMP-binding capability have also been discovered ^8–11^. Bacterial genomes encode a large number of PilZ domain-containing proteins that feature different protein length and domain composition ^1, 12^. Currently, there is more than 28,000 PilZ domain-containing proteins listed in the InterPro protein database ^13^. However, only a small number of PilZ proteins have been examined experimentally as of today. Even for the model microorganisms (e.g. *Pseudomonas aeruginosa*, *Vibrio cholerae*, and *Xanthomonas campestris pv. campestris*) that have been under intensive investigation to understand the molecular mechanisms of c-di-GMP signaling, the cellular function for most of the PilZ proteins remain unknown. As PilZ proteins usually share weak sequence homology, it remains a challenge to deduce the function of functionally unknown PilZ proteins solely based on the sequence-function relationship.

Newly discovered proteins are routinely annotated with predicted function based on the function of experimentally characterized homologs ^14^. This approach assumes that the sequence of a protein dictates its structure and function, and that proteins with high sequence similarity are likely to share the same or similar functions. By taking this approach, the function of a few experimentally characterized proteins can be extrapolated to a large number of homologs or orthologs. Many computational methods have been developed to predict protein function based on sequence homology and sequence-function relationship. Among the computational methods, phylogenetic relationship analysis is often used for sequence-function relationship analysis. However, the construction of phylogenetic trees consisting of large sequence sets is computationally intensive and interpretation is difficult as the trees increase in size and complexity ^15^. To address this problem, Babbitt and Gerlt et al developed the Sequence Similarity Networking (SSN) tool for the visualization of complex relationships and functional trends across protein sequences ^15–16^. In SSN analysis, the relationship between proteins is measured by independent pairwise alignments between sequences and the entire context is visualized in the form of a network. In an SSN, proteins are represented by nodes and are connected to other nodes by edges or connecting lines that represent the sequence similarity of the nodes ^16^. Only nodes that share pairwise sequence alignments greater than a user-specified value are connected by edges to create a network that reveals potential functional relationships among the proteins. Generating SSNs is less computationally demanding than phylogenetic tree analysis and a large amount of sequence data can be displayed and analyzed. The display of sequence relationships in a two-dimensional topology is also more visually intuitive and thereby has attracted an increasing number of users. Today, SSN is mainly used by enzymologists to explore sequence similarity and function diversification through the segregation of diverse superfamilies of enzymes into putative isofunctional groups ^16^. The analysis has led to the discovery of new catalytic reactions or substrate specificity within enzyme superfamilies ^17–20^.

With the main objective of exploring the sequence and function space of PilZ domain proteins, here we apply the SSN tool to single and di-domain PilZ proteins. SSNs were constructed for single- and di-domain PilZ proteins separately with experimentally characterized PilZ proteins (“reference nodes”) mapped onto the networks to aid the generation of isofunctional clusters. We also generated sequence logos to identify conserved functional residues or motifs to corroborate the formation of isofunctional clusters. The study reveals a great degree of sequence variation and likely function diversification in PilZ proteins. Considering that only about a dozen of PilZ proteins have been experimentally characterized, we hope that the SSNs will provide an overarching view of the scope of the functional diversification of PilZ proteins, and generate a useful framework for formulating hypotheses about the cellular function of uncharacterized PilZ proteins.

## Results

### Classification of PilZ proteins into single, di- and multi-domain proteins

The core of PilZ domain features a β-barrel of 68-74 residues ^5–6, 21–22^. The canonical c-di-GMP-binding PilZ domain is also characterized by a flexible N-terminal loop (12-18 amino acids) that harbors the key RxxxR motif involved in direct c-di-GMP binding. We identified and collected a total of 28,223 sequences of PilZ domain protein sequences from the InterPro database, with most of the sequences belonging to the IPR009875 and IPR031800 domain families. Several non-canonical PilZ domain proteins that lack the RxxxR and (D/N)xSxxG motifs have been reported ^8, 10, 23^. These non-canonical PilZ proteins, which cannot be found in either the IPR009875 or IPR031800 protein families, are also included in our analysis.

The sequence-length histogram generated for the 28,223 PilZ proteins shows that 13,476 of them contain 80 to 140 residues (Figure 1). We classified this group of proteins as single-domain PilZ proteins considering that they only contain a core PilZ domain, an N-terminal loop and a short C-terminal motif. The variation in protein length in the single-domain PilZ proteins is mainly due to the differences in the C-terminal region, with secondary structure analysis suggesting that the C-terminal region exhibits great structural diversity by adopting helical, strand or coil secondary structure. The second-largest group of PilZ proteins (9,053 in total) contains 140 – 280 residues; and we classified them as di-domain PilZ proteins because they possess an additional C- or N-terminal protein domain. YcgR_N_ domain ^2, 6, 24^ is by far the most abundant domain found in the di-domain PilZ proteins. Besides the YcgR_N_ domain, the phosphoreceiver (REC) domain, DNA-binding domain, GGDEF domain or a second PilZ domain are also commonly found in the di-domain PilZ proteins.

**Figure 1.**
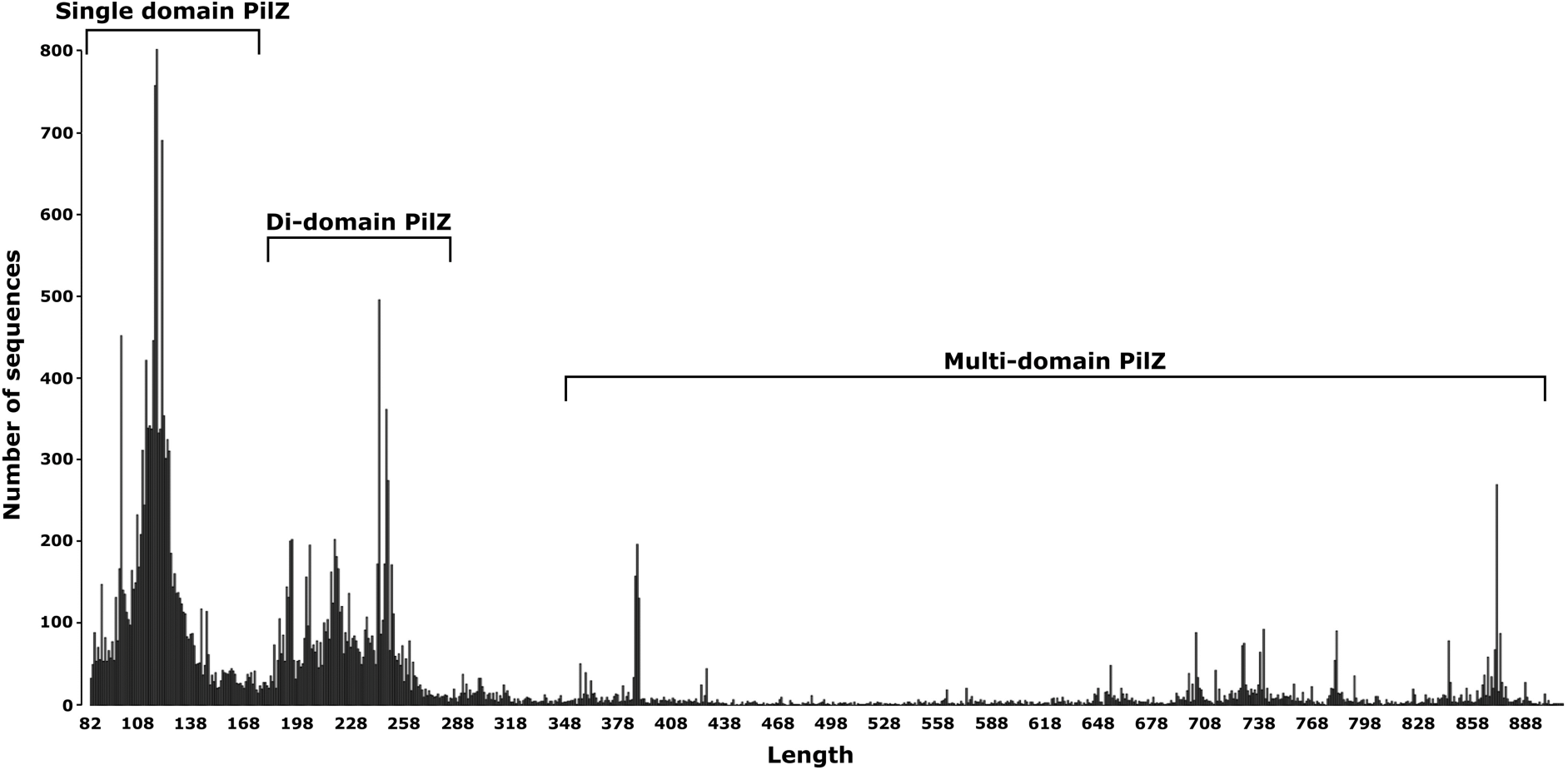
Length histogram for the 28,223 proteins from the PilZ protein superfamily. The histogram shows the number of sequences as a function of sequence length. The PilZ protein superfamily is grouped into single-, di- and multi-domain PilZ proteins families based on the sequence length.

Apart from the single and di-domain PilZ proteins, the rest of PilZ proteins feature much longer protein sequences (>300 amino acids) and are considered to be multi-domain proteins based on InterPro and Pfam domain analysis. The multi-domain PilZ proteins contain a variety of protein domains, with the most common domains predicted to be glycosyltransferase, GGDEF, EAL, methyl-accepting chemoreceptor and other sensory protein domains. The variation in protein length and domain composition indicate the PilZ domain has been harnessed to regulate different cellular functions through gene fusion. The biological function of many multi-domain PilZ proteins can be inferred with high confidence based on domain annotation and characterized homologs. For example, the largest group of multiple PilZ domain proteins, which has an average sequence length of 880 residues, contains a glycosyltransferase domain and shares high similarity with the cellulose synthase subunit of *Escherichia coli*, *Rhodobacter sphaeroides* and *Gluconacetobacter xylinum* ^22, 25–26^. Those proteins are likely to be involved in the production of cellulose and other types of extracellular polysaccharides (EPS). Likewise, the ∼200 multi-domain PilZ proteins that share high sequence homology with Alg44 (388 aa), a multi-domain PilZ protein involved in alginate biosynthesis in *Pseudomonas aeruginosa* ^27–28^, are likely to be involved in the production of EPS as well. The studies described below will focus on the analysis of the functionally more diverse and more challenging single and di-domain PilZ proteins, but not the multi-domain PilZ proteins.

### Overview of the SSN for single-domain PilZ proteins

We generated a series of SSNs for the 13,476 single-domain PilZ proteins using a series of alignment scores. We then located the reference nodes, i.e., the PilZ proteins with a known structure or biological function, to find out whether the known function homologs are grouped in the same clusters. We found that by using the alignment score of 20 (∼35% identity) and a thresholding BLAST E-value of 1 × 10^-20^, the proteins are potentially sorted into isofunctional clusters that contain function orthologs/homologs as suggested by the clustering of the reference nodes ^16^. The final representative network consists of 8,025 meta-nodes distributed in clusters of various sizes (Figure 2). Note that the proteins with sequence identity greater than 90% are merged into one single meta-node in the representative SSN. The observation of a large number of clusters is consistent with the low sequence similarity shared by the single-domain PilZ domains. Some of the clusters appear compact with short edges connecting tightly packed nodes; while others feature more scattered nodes that can be further grouped into subclusters. As the nodes are connected by edges whose length reflects the sequence similarity shared by the proteins ^16^, the lack of compactness and presence of subclusters may reflect a further function divergence within some of the clusters. The nodes in the SSN are colored according to the taxonomical order of the bacteria from which the sequences come from. Clusters consisting of sequences originated from various taxonomical order signify a conserved role of the PilZ homologs across bacterial species; whereas clusters consisting sequences exclusively from a single order or genus indicate an order- and genus-specific role of the PilZ proteins.

**Figure 2.**
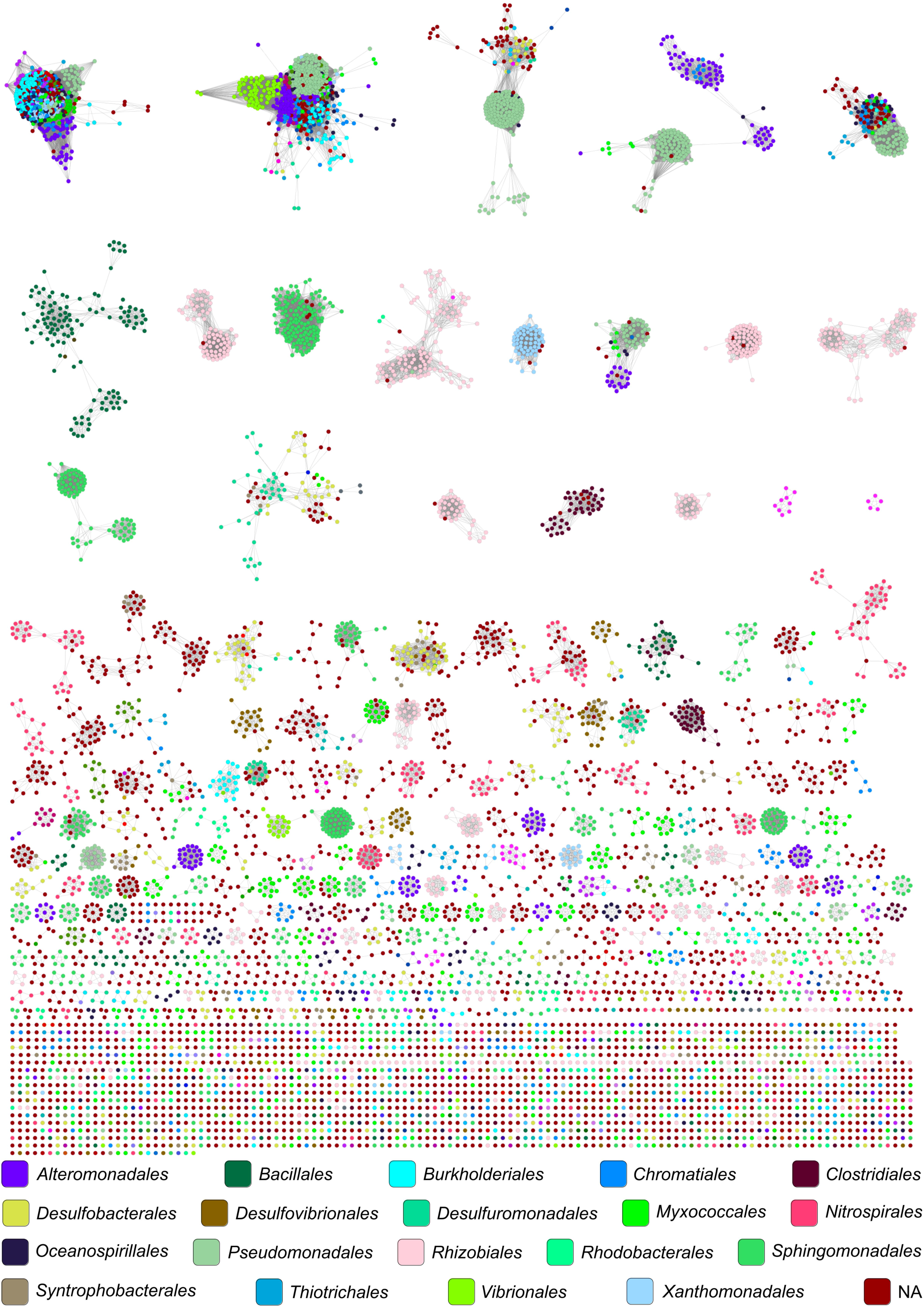
The SSN provides a global view of sequence relationships in single-domain PilZ proteins. The SSN contains 13,476 sequences represented by 8,028 meta-nodes filtered at 90% sequence identity. The nodes are colored based on the taxonomic order of the bacteria from which the protein is from. Edges or lines between nodes are shown if the least significant pairwise sequence similarity score between the representative sequences of two nodes is high than the threshold (BLAST E-value less than 1 × 10^-20^).

The 18 clusters with the largest number of proteins account for 48% of the total single-domain PilZ proteins. We arranged the 18 clusters according to the number of sequences they contain and mapped experimentally characterized PilZ domains onto the SSN to find out how they are related to other functionally unknown proteins (Figure 3). The two small clusters that contain DgrA and DgrB, two functionally known PilZ proteins from *Caulobacter crescentus*, are also included in Figure 3. Notably, the five single-domain PilZ proteins (HapZ, MapZ, PA0012, PA2960, PA4324) from *P. aeruginosa* are found in five different clusters. Likewise, the single-domain proteins from several other bacterial species (e.g. *V. cholerae* and *X. campestris pv. campestris*) are also distributed in different clusters. Below we will focus our analysis on the clusters that contain experimentally characterized PilZ protein to demonstrate that some of the clusters in the SSN are likely isofunctional clusters, and thus, the SSN may have captured the scope of function divergence for the single-domain PilZ proteins.

**Figure 3.**
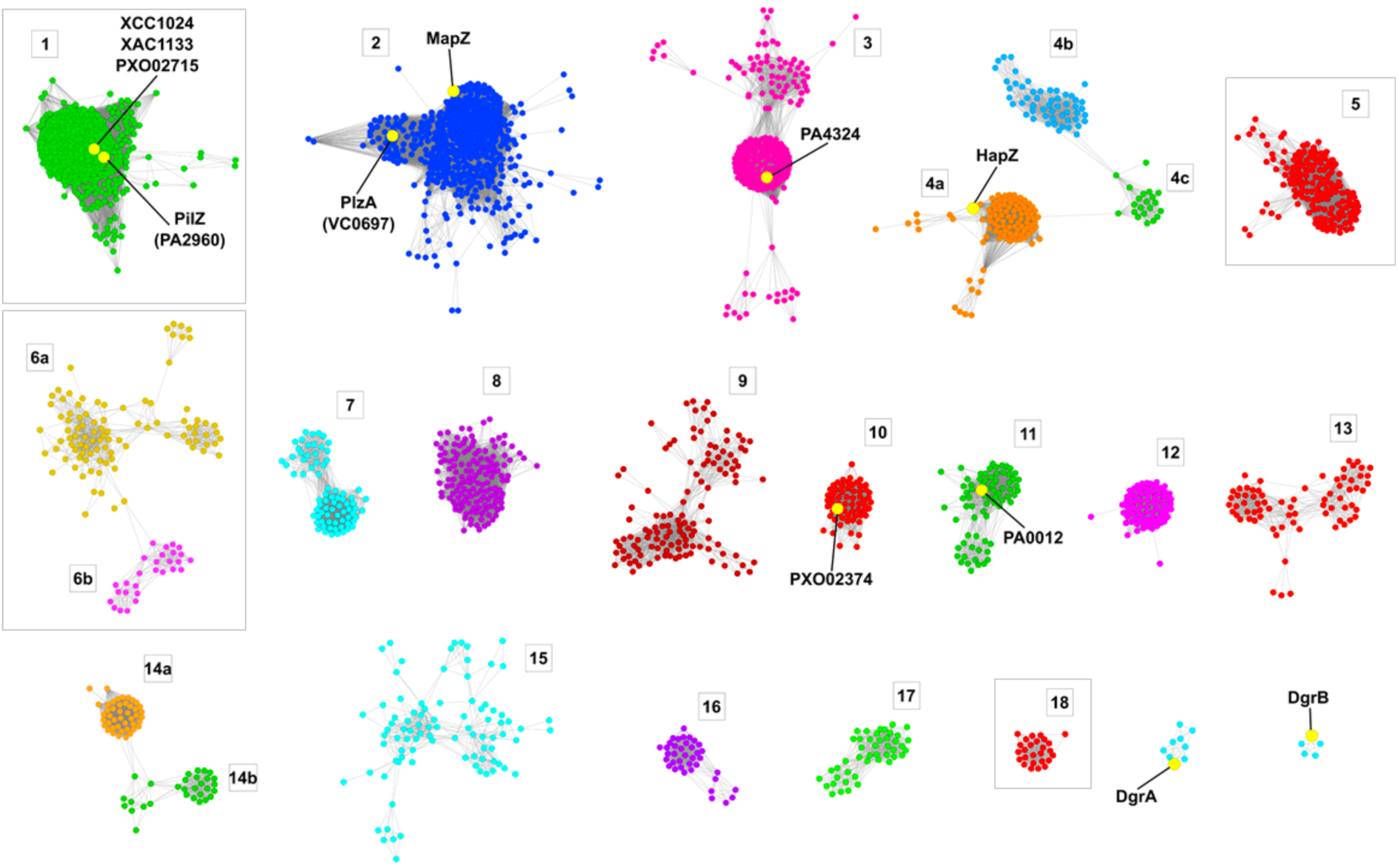
Eighteen large clusters from the single-domain PilZ protein SSN that contain more than 75 sequences. Two small clusters that contain the characterized DgrA and DgrB are also shown here. The reference nodes or proteins documented in the literature are labeled. The nodes are colored to indicate sub-clustering within the clusters. The clusters that feature defective RXXXR or D/NXSXXG motif are shown in the box.

### SSN analysis reveals an unexpectedly large number of non-canonical PilZ proteins that lack c-di-GMP-binding motifs

Canonical c-di-GMP-binding PilZ domains are characterized by two conserved motifs: the arginine-rich motif (RxxxR) from the N-terminal loop and the (D/N)xSxxG motif from the β-strands. Inspection of the sequence logo for the 18 largest clusters reveals the two motifs are conserved in most clusters except for clusters 1, 5, 6 and 18 (Fig. 4), which indicates that the proteins from the four clusters are likely to have a low or negligible c-di-GMP binding affinity.

As the largest cluster, cluster 1 consists of 1,779 sequences from at least 195 genera and accounts for 13% of all single-domain PilZ proteins (Figure 5A). The relative compactness of this cluster reflects the high sequence similarity shared by the proteins. The cluster contains four proteins that are documented in the literature: PilZ (PA2960, *P. aeruginosa*), XCC1028 (also by the name XCC1024 in some publications) (*Xanthomonas campestris pv. campestris*), XAC1133 (*Xanthomonas citri subsp. citri*) and PXO_02715 (*Xanthomonas oryzae pv. oryzae*). The three proteins XCC1024, XAC1133 and PXO_02715 share >90% sequence identity and hence merge into a single node. Binding assays showed that the four proteins do not exhibit noticeable c-di-GMP binding affinity ^8–10, 28–29^. It has been shown that XCC1024 and XAC1133 can form complex with FimX and PilB orthologs to regulate Type IV Pilus biogenesis ^8, 10^. Despite the high sequence similarity shared XCC1024 and XCA1133, PilZ (PA2960) does not interact with FimX in *P. aeruginosa* due to the replacement of several key residues in *P. aeruginosa* FimX ^10^; whereas PilB and FimX interact directly with each other without involving PilZ (PA2960)^10^. PXO_02715 modulates the expression of T3SS genes and bacterial virulence by interacting with the FimX homolog Flip in *X. oryzae pv. Oryzae* ^30^. The crystal structures of XCC1024 and XAC1133 in complex with their protein partner (i.e. the EAL domain of FimX) have been determined to reveal the regions and residues important for protein-protein interaction. The residues in XAC1133 that mediates the interaction with FimX are conserved for the proteins from this cluster (Figure 5B) ^8^. Hence, the PilZ proteins from this cluster are likely to function as adaptors to mediate the protein-protein interaction in pili biogenesis.

**Figure 4.**
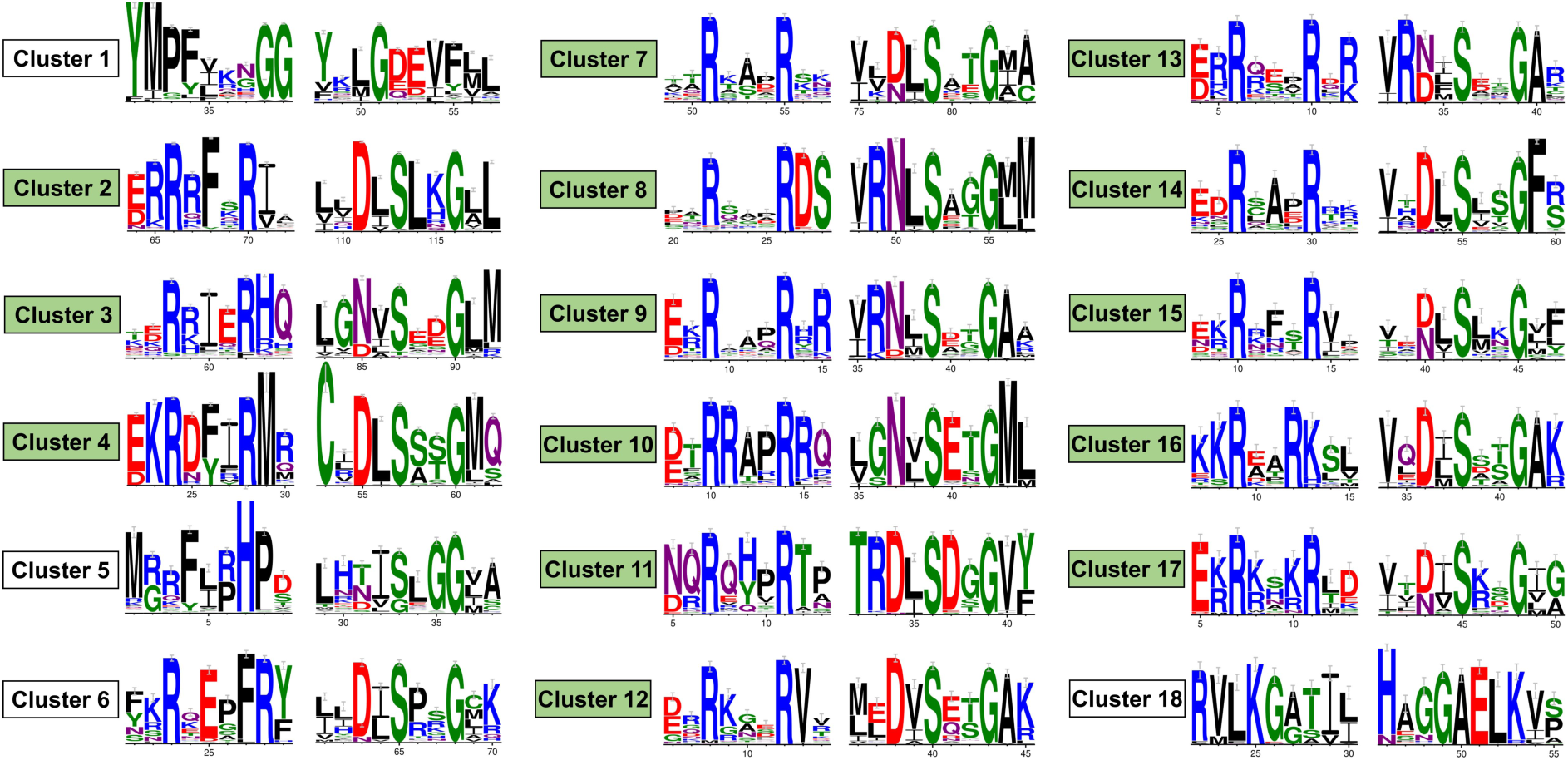
Partial sequence logs showing sequence variation within and near the two RxxxR and N/DxSxxG motifs in the 18 large single-domain PilZ proteins. The two motifs are absent in the proteins from clusters 1, 5 and 18. The RxxxR motif is replaced by an RxxxFR motif in cluster 6.

**Figure 5.**
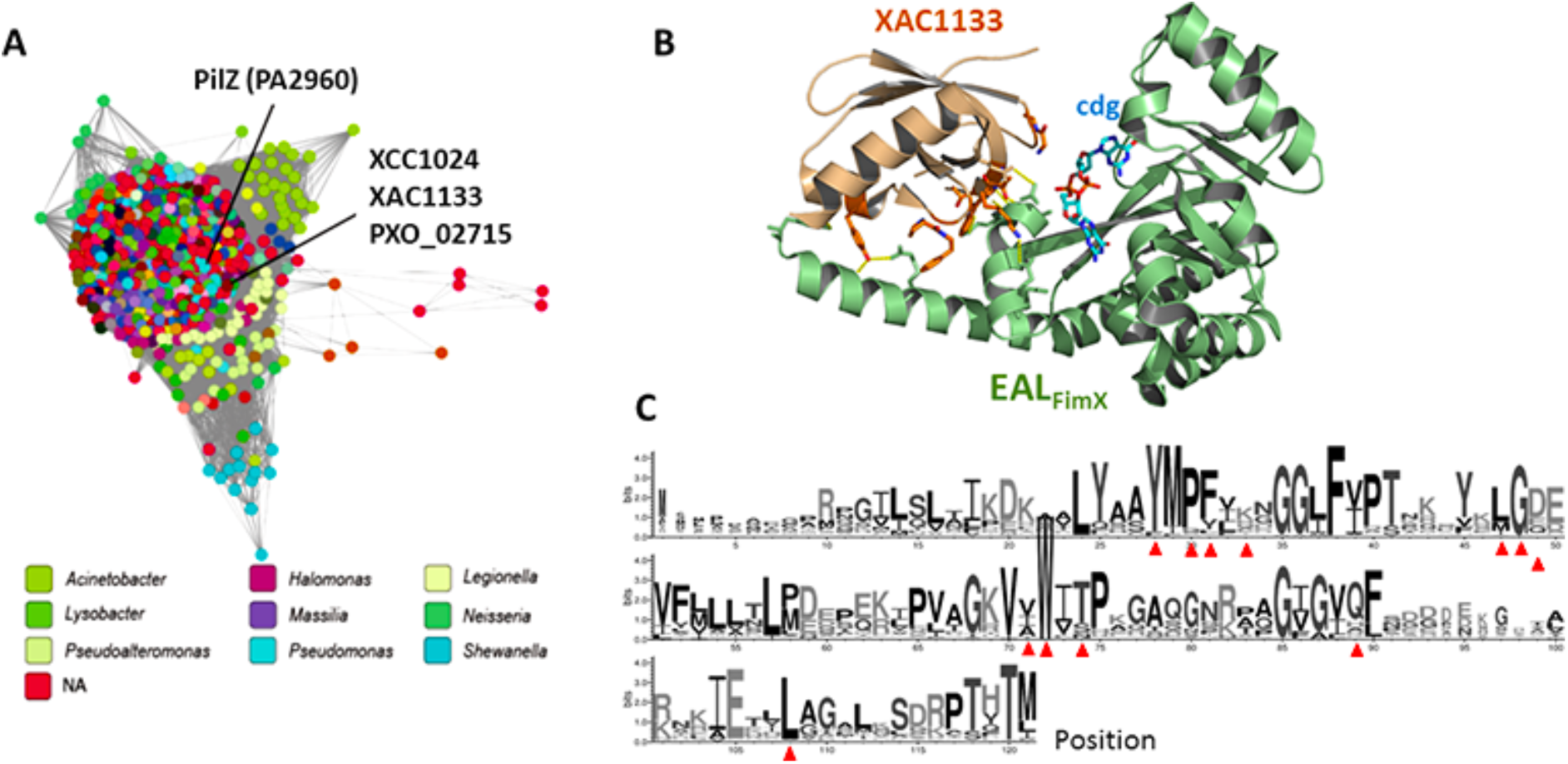
Cluster 1 contains a group of widely distributed non-canonical PilZ proteins. A. The representative SSN for cluster 1. The nodes colored according to the genus of the bacteria from which the sequences are from. The four reference nodes (i.e., characterized proteins) are labeled and the three reference proteins that share more than 90% sequence identity merge into one single node. B. The crystal structure of the XAC1133/EAL_FimX_ complex (PDB: 4FOU) with the residues involved in the direct binding of EAL_FimX_ labeled and shown as sticks. C-di-GMP bound by EAL_FimX_ is also shown in sticks and labelled as cdg. C. Sequence logo for the PilZ proteins from cluster 1. The residues that are directly involved in binding EAL_FimX_ are indicated by the red triangles ^66–67^.

The proteins from clusters 5 and 18 also lack the two c-di-GMP-binding motifs and are predicted to lack c-di-GMP binding ability. Cluster 5 contains 304 sequences from a variety of genera while cluster 18 contains 77 sequences from the genera of *Mesorhizobium* and *Aminobacter*. None of the proteins from the two clusters are experimentally characterized. Meanwhile, cluster 6 contains 239 sequences from Gram-positive bacteria exclusively. The proteins from this cluster are distinct from the canonical PilZ proteins by featuring an unusual RxxxFR motif, in comparison to the canonical RxxxR motif. As the RxxxR motif seems to be essential for c-di-GMP binding, the PilZ proteins from this cluster may have also lost the ability to bind c-di-GMP.

### A large group of single-domain PilZ proteins predicted to interact with chemotaxis methyltransferases

Cluster 2 contains a total of 1,173 sequences that account for 9% of the single-domain PilZ proteins. Cluster 2 is the second-largest cluster in the SSN and is a relatively compact cluster with most proteins sharing high sequence similarity (Figure 6A). Cluster 2 represents a group of PilZ proteins found in more than 97 genera of proteobacteria, with more than half of them belonging to *Pseudomonas* and *Vibrio* (Figure 6A). Considering that all the proteins from this cluster contain the RxxxR and (D/N)xSxxG motifs for c-di-GMP binding (Figure 6B), they are likely to function as c-di-GMP-binding receptors or adaptors.

**Figure 6.**
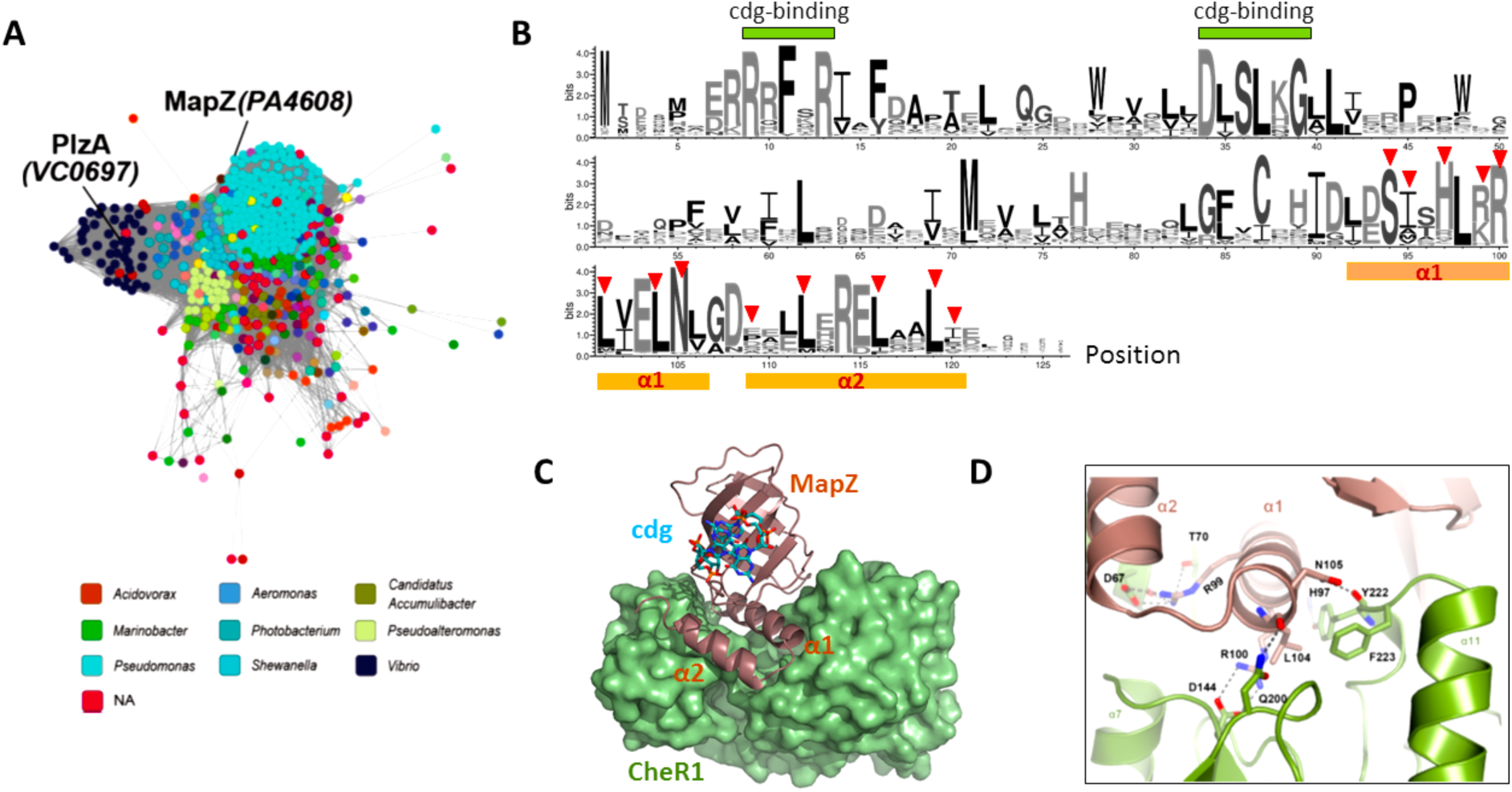
Cluster 2 contains a group of MapZ homologs/orthologs predicted to interact with chemotaxis methyltransferases. A. Cluster 2 with the nodes colored according to the genus of the bacteria from which the sequences are from. The two characterized proteins are labeled. B. Sequence logo for the proteins from Cluster 2. The residues that are involved in the direct binding of CheR1 are indicated by the red triangles. C. The crystal structure of the MapZ/CheR1/c-di-GMP ternary complex (PDB: 2L74) shows the protein-protein interaction is mainly mediated by α1 and α2 of MapZ. D. Specific protein-protein interaction between CheR1 and the conserved residues from the helix α1 of MapZ. The R99 residue is essential for CheR1-MapZ interaction as the R99A mutation reduced the binding affinity to below the detection limit ^7^.

Within this cluster is MapZ, a single-domain PilZ protein from *P. aeruginosa* PAO1. MapZ is one of the best characterized PilZ proteins with its structure, interacting partner and physiological function elucidated ^5, 31–34^. In vitro and in vivo studies revealed that MapZ binds c-di-GMP with a *K*_D_ in the range of 6-9 µM ^5, 32^, and that it interacts with the chemotaxis methyltransferase CheR1 at elevated c-di-GMP concentrations. Crystallographic studies on the MapZ/CheR1/c-di-GMP ternary complex identified that the two C-terminal helices α1 (residues 92-106aa) and α2 (residues 109-121) of MapZ are the key motifs mediating MapZ-CheR1 interaction (Figure 6C) ^7, 33^. In particular, the helix α1 plays an essential role in mediating the MapZ-CheR1 interaction by occupying the central cleft of CheR1. Several residues from α1, including R99, R100, S96, H97 and N105 interact with various CheR1 residues through hydrogen bonds and salt bridges (Figure 6D); while the side chain of several other residues, including L92, I95, L101, L104, R99, R100 and E103, interact with various CheR1 residues through hydrophobic interaction. Several non-polar residues from the helix α2 are also involved in the interaction with CheR1 through hydrophobic interactions. Importantly, the sequence logo shows that most of the key CheR1-interacting residues from α1 and α2 are strongly conserved (Figure 6B). Conservation of the functionally important residues strongly suggests that the proteins in cluster 2 are likely to regulate chemotaxis by interacting with CheR1 homologs. Another PilZ proteins from this cluster that was documented in the literature is PlzA, a PilZ protein from *V. cholerae* with an unknown cellular role ^35^. The clustering of PlzA with MapZ, together with the conserved CheR1-binding residues, hints a role of PlzA in the regulation of chemotaxis and bacterial motility.

### Single-domain PilZ proteins predicted to interact with phosphor-receiver (REC) domains

Cluster 4 contains 593 sequences and seems to contain a few subclusters. The two motifs RxxxR and D/NxSxxG for c-di-GMP binding are conserved in this cluster of proteins, indicating that the proteins are capable of binding c-di-GMP.

Cluster 4a contains 436 sequences with the majority of them from the genus *Pseudomonas*. The majority of the sequences in this subcluster share high homology as reflected by the compactness of the subcluster. The only protein from this cluster with known biological function is the *P. aeruginosa* PAO1 protein HapZ (previously PA2799). HapZ functions as a c-di-GMP binding adaptor protein to regulate two-component signaling pathways ^36^. Specifically, HapZ interacts with the REC domain of the histidine kinase SagS at elevated c-di-GMP concentrations to control the transfer of phosphate group from SagS to its downstream proteins. The high sequence similarity shared by the sequences from subcluster 4a prompts us to propose that the other uncharacterized proteins from the subcluster also interact with REC domains. As REC domain is a widely distributed signaling domain, we surmise that the HapZ homolog/orthologs may interact with REC domain-containing histidine kinases, response receivers or other signaling proteins. Besides, the sequence logo also reveals several conserved residues (C36, E54, V79, R81, F/L73) that are unique to this cluster of proteins and are potentially involved in the interaction with their protein partners.

### Other major clusters of c-di-GMP-binding single-domain PilZ proteins feature distinct C-terminal motifs and conserved residues

Several other clusters from the SSN also contain one or more proteins that have been documented in the literature. The third-largest cluster (cluster 3) consists of proteins from various genera, including the protein PA4324 from *P. aeruginosa*. The physiological function of this group of widely occurred proteins remains unknown at this moment. We noticed that this group of proteins contains a distinct hydrophobic C-terminal segment with several invariant non-polar residues (L106, Y109, F110, F112 and P114). As such hydrophobic C-terminal region is unique for single-domain PilZ proteins; we speculate that the hydrophobic residues may be involved in the direct binding of their protein partners. Cluster 10 is a compact cluster comprised of proteins mainly from the *Xanthomonas* genus. The cluster contains the *X. oryzae pv. oryzae* protein PXO_02374, a protein that regulates bacterial virulence by controlling T3SS gene expression ^37^. Cluster 11 consists of 173 proteins, including the PilZ protein PA0012 that regulates bacterial virulence in *P. aeruginosa* (Cheang et al, to be published). C-di-GMP signaling proteins have been intensively studied in the modal organism *Caulobacter crescentus* ^38–41^. Two dozens of PilZ proteins from the *Caulobacter* genus form two distinct small clusters separated from other PilZ proteins in the network with two small clusters contain proteins exclusively from the *Caulobacter* genus. One cluster contains DgrA and the other contains DgrB. Both DgrA and DgrB are known to suppress bacterial motility at elevated levels of c-di-GMP ^40^.

The physiological function of the single-domain PilZ proteins from the rest of the clusters remain completely unknown at this moment. Some of the clusters contain proteins that are exclusively or mostly from a single genus or order, indicating that the proteins were evolved to regulate genus- or order-specific cellular function. For example, cluster 7 consists of 218 proteins that are mostly from the *Rhziobium* genus and cluster 12 consists of 170 proteins that are mostly from the *Bradyrhizobium* genus. Some other clusters, such as clusters 14 and 15, contain proteins from a diversity of genera to indicate a broader distribution of the PilZ-mediated mechanism.

### Overview of the SSN for di-domain PilZ proteins

The SSN for the 9,053 di-domain PilZ proteins was generated using a 40% sequence similarity cut-off (E-Value of 10^-25^). The sequences were grouped into clusters of various sizes by using an alignment score of 25. The alignment score was chosen because it potentially segregates the nodes into isofunctional clusters according to the clustering of reference nodes. The clusters are arranged from the largest to the smallest in the final SSN shown in Figure 7. Sequence logos were generated for the 15 largest clusters that contain more than 75 sequences to identify conserved residues and motifs. We colored the nodes of the SSN according to the taxonomical order to generate a snapshot of the relative distribution and abundance of the di-domain PilZ proteins (Figure 7). Similar to the single-domain PilZ SSN, the SSN is dominated by proteins from proteobacteria (α, β, γ and Δ-*proteobacteria*) and firmicutes (mainly *Bacilli*, *Clostridia* and *Nitrospira*).

**Figure 7.**
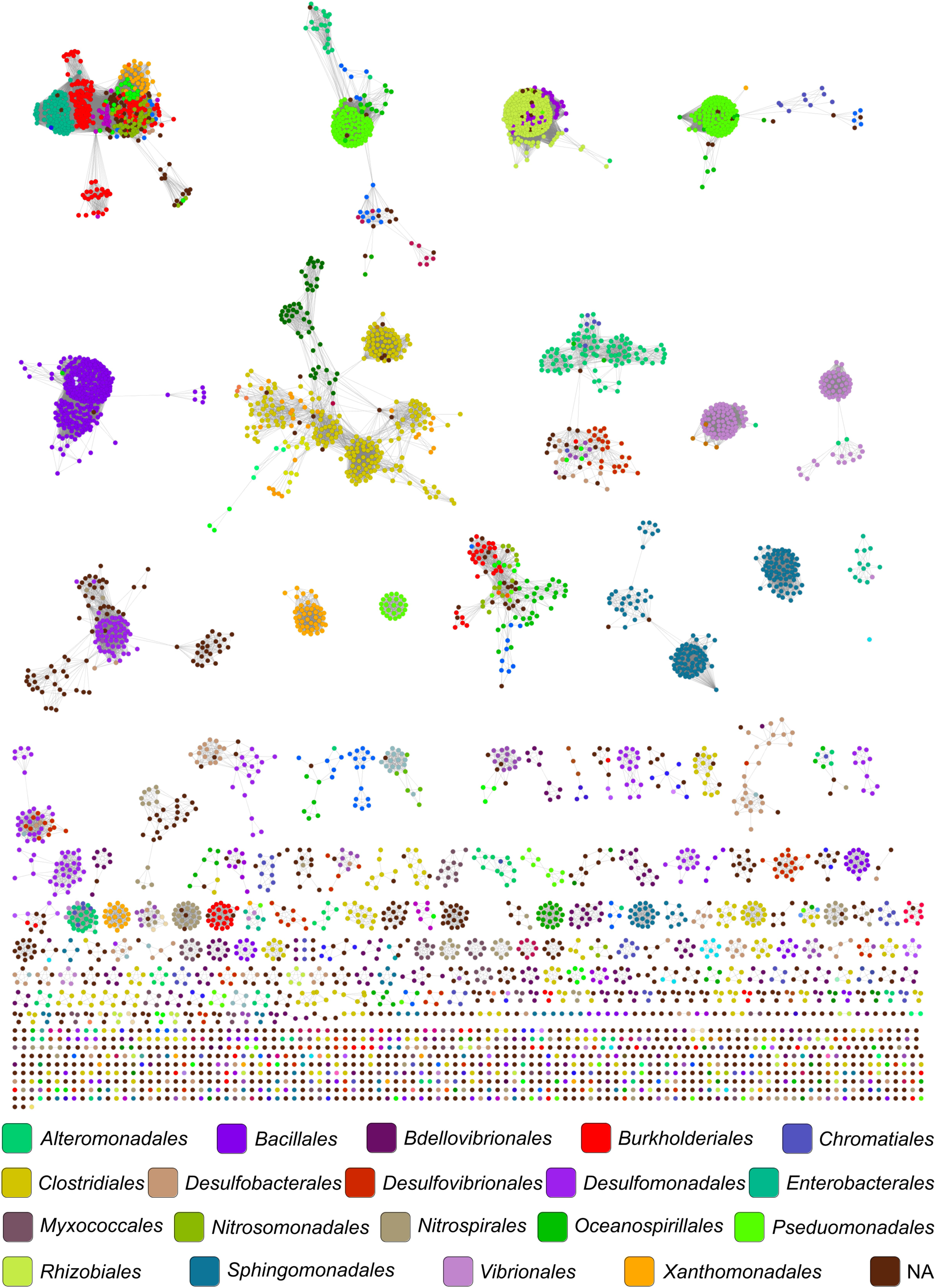
The SSN provides a global view of sequence relationships in single-domain PilZ proteins. The SSN contains 9,053 sequences represented by 4,963 meta-nodes filtered at 90% sequence identity. The nodes are colored based on the order of the bacteria from which the proteins are from. Edges or lines between nodes are shown if the least significant pairwise sequence similarity score between the representative sequences of two nodes is high than the threshold (BLAST E-value less than 1 × 10^-25^).

Only a handful of di-domain PilZ proteins have been documented in the literature. We identified these proteins in the SSN and use them as reference nodes to guide the segregation of nodes into putative isofunctional clusters (Figure 8). One of the first characterized di-domain PilZ proteins is the YcgR protein from *E. coli* ^2, 24, 42–43^. A large number of di-domain PilZ proteins are predicted to be YcgR–like proteins that feature an N-terminal YcgR_N_-like domain and a C-terminal c-di-GM-binding PilZ domain. In the SSN, YcgR-like proteins constitute 38% of the total proteins and many are found in large clusters (Clusters 1, 4, 5, 6, 7, 8, 9 and 13). PilZ domain is also fused to a variety of other non-YcgR_N_ domains to form non-YcgR-type di-domain proteins. The non-YcgR type proteins can be found in the large clusters 2, 3, 10, 11, 12, 14, 15 as well as many smaller clusters, including the two clusters that contain MrkH and BB0733.

**Figure 8.**
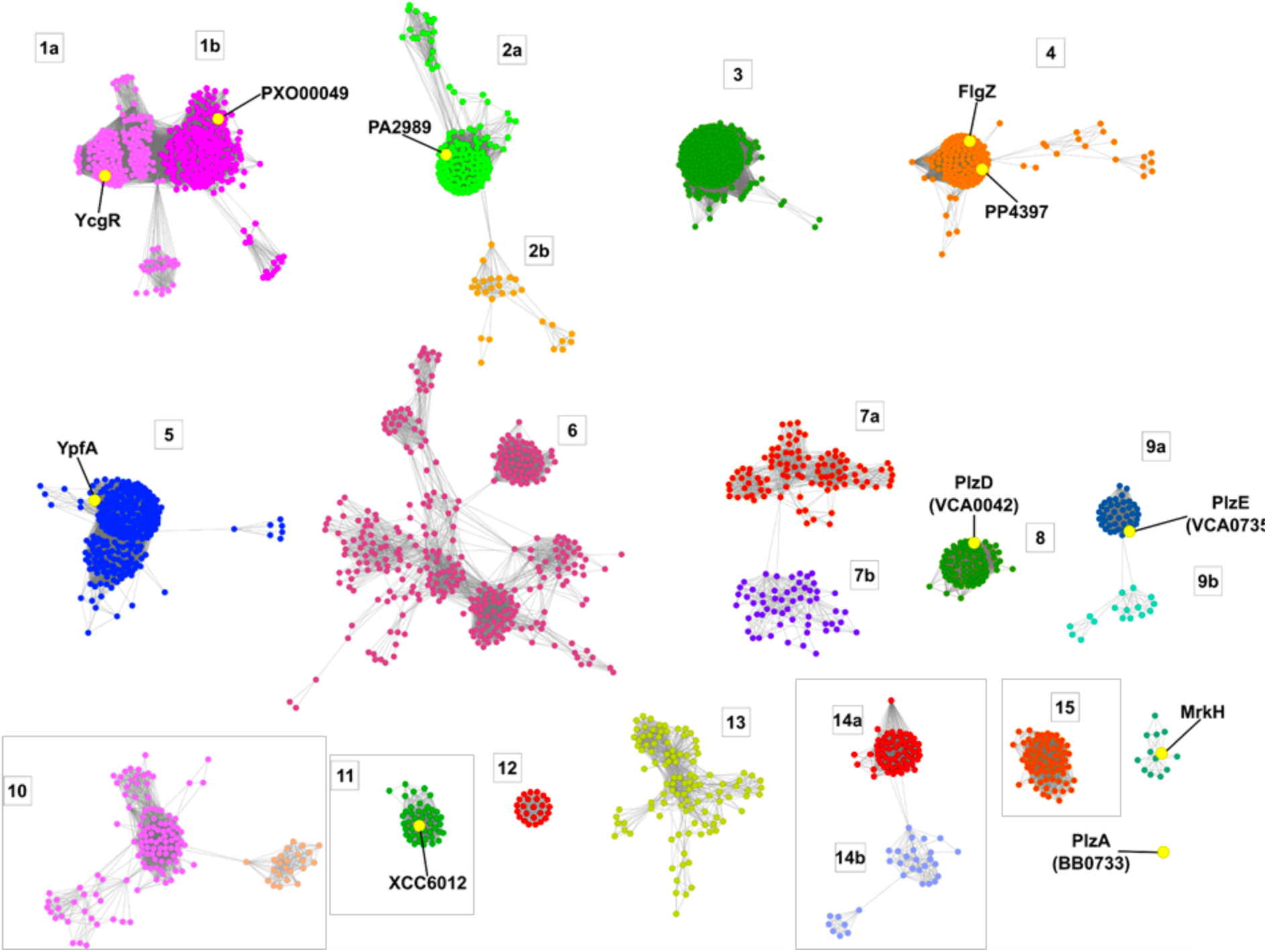
Fifteen large clusters from the SSN of single-domain PilZ proteins that contain more than 75 nodes. Two small clusters that contain the characterized MrkH and PlzA (BB0733) are also shown here. The reference nodes or proteins documented in the literature are labeled. The nodes are colored to indicate sub-clustering within the clusters. The clusters that feature defective RXXXR or D/NXSXXG motif are shown in the box.

As di-domain PilZ proteins also use the RxxxR and D/NxSxxG motifs for binding c-di-GMP, we inspected the sequences to identify the proteins that may be defective in c-di-GMP binding. We found that most of the di-domain proteins contain the two motifs with the exception of proteins from four clusters. The most notable one is cluster 11, which features protein sequences with the two motifs completely missing (Figure 9). The RxxxR motif is replaced by an RxxxxxR motif for the majority of proteins in cluster 10, and one of the R residues in the RxxxR motif is missing in a minority of proteins in clusters 14 and 15 (Figure 9). We also noted that the D/NxSxxG motif in the proteins of cluster 2 is replaced by an xxSEGG motif. The alteration of the D/XxSxxG motif in this cluster of proteins does not seem to affect c-di-GMP binding as we discuss further below.

**Figure 9.**
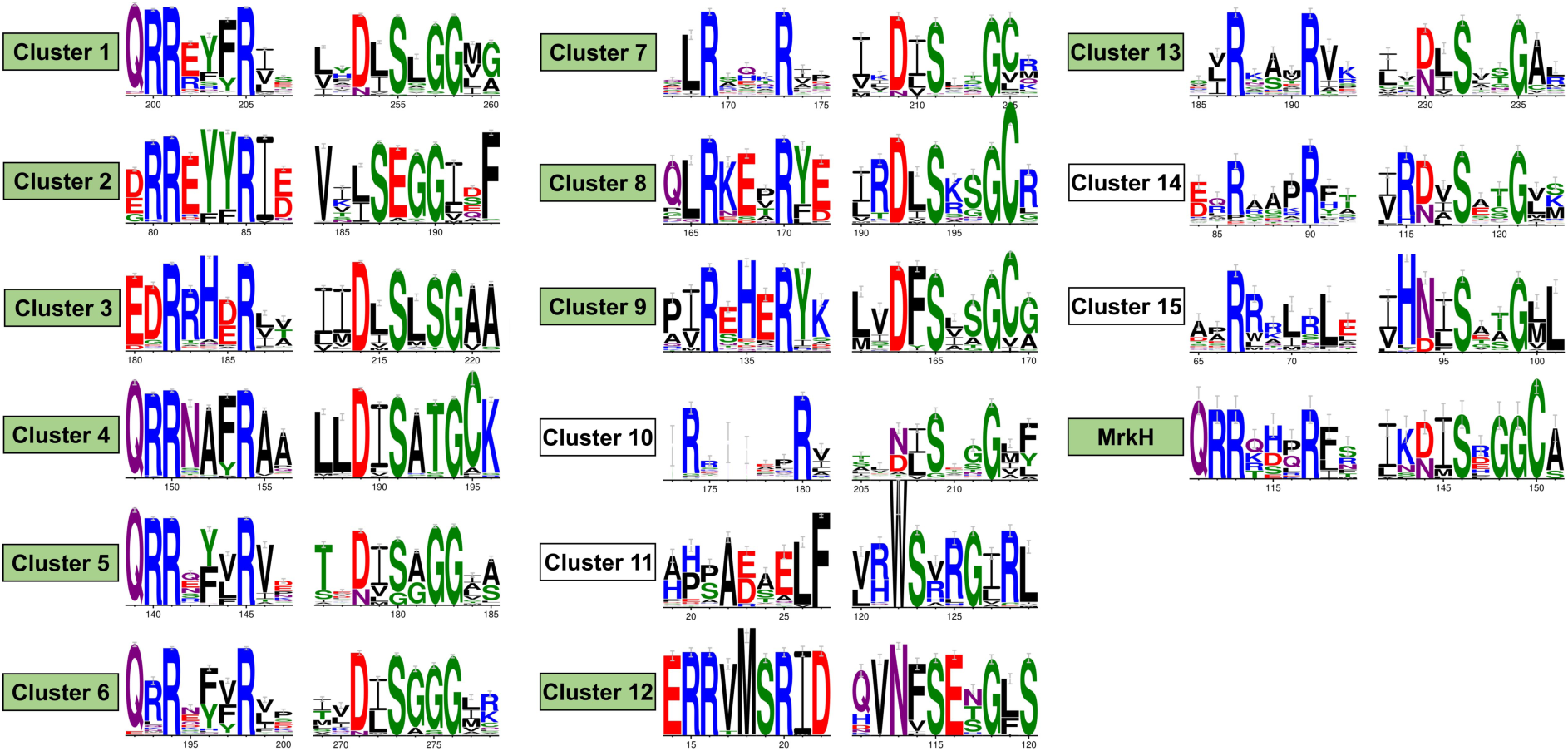
Sequence variation within and near the two RxxxR and N/DxSxxG motifs in the 15 largest di-domain PilZ proteins and the MrkH cluster. The two motifs are absent in the proteins from clusters 11. A portion of proteins from clusters 5 and 18 do not contain the intact RxxxR motif. The RxxxR motif is replaced by an RxxxxxR motif in cluster 10.

### YcgR-like PilZ proteins exhibit great sequence and function divergence

Cluster 1, which consists of 1,398 sequences exclusively from proteobacteria, is the largest in the SSN. The cluster consists of two major subclusters (1a and 1b). Subcluster 1a contains YcgR, the *E. coli* protein that interacts with the flagellar switch-complex protein FliM/FliG to act as a “backstop brake”^42, 44^. It was proposed that YcgR interacts with FliM and FliG using both the PilZ domain and the YcgR_N_ domain ^44^. Although the crystal structure of YcgR has not been determined, the residues important for FliM and FliG interaction were identified based on a structural model ^24^. The sequence logo generated for the proteins from subcluster 1a suggests a high degree of conservation of the residues involved in binding FliG/FliM (Figure 10A), which hints that this group of proteins is potentially true functional orthologs/homologs of YcgR. In contrast, cluster 1b contains a group of proteins that lack the key residues for FliG and FliM binding. One of the proteins from cluster 1b, PXO_00049 from the plant pathogen *X. oryzae pv. Oryzae*, was reported to mediate sliding motility, type III secretion system (T3SS) gene expression and bacterial virulence ^37^. The interacting partner of PXO_00049 remains unknown, and the lack of FliG/FliM binding residues suggests that it may interact with a different protein other than FliM/FliG.

**Figure 10.**
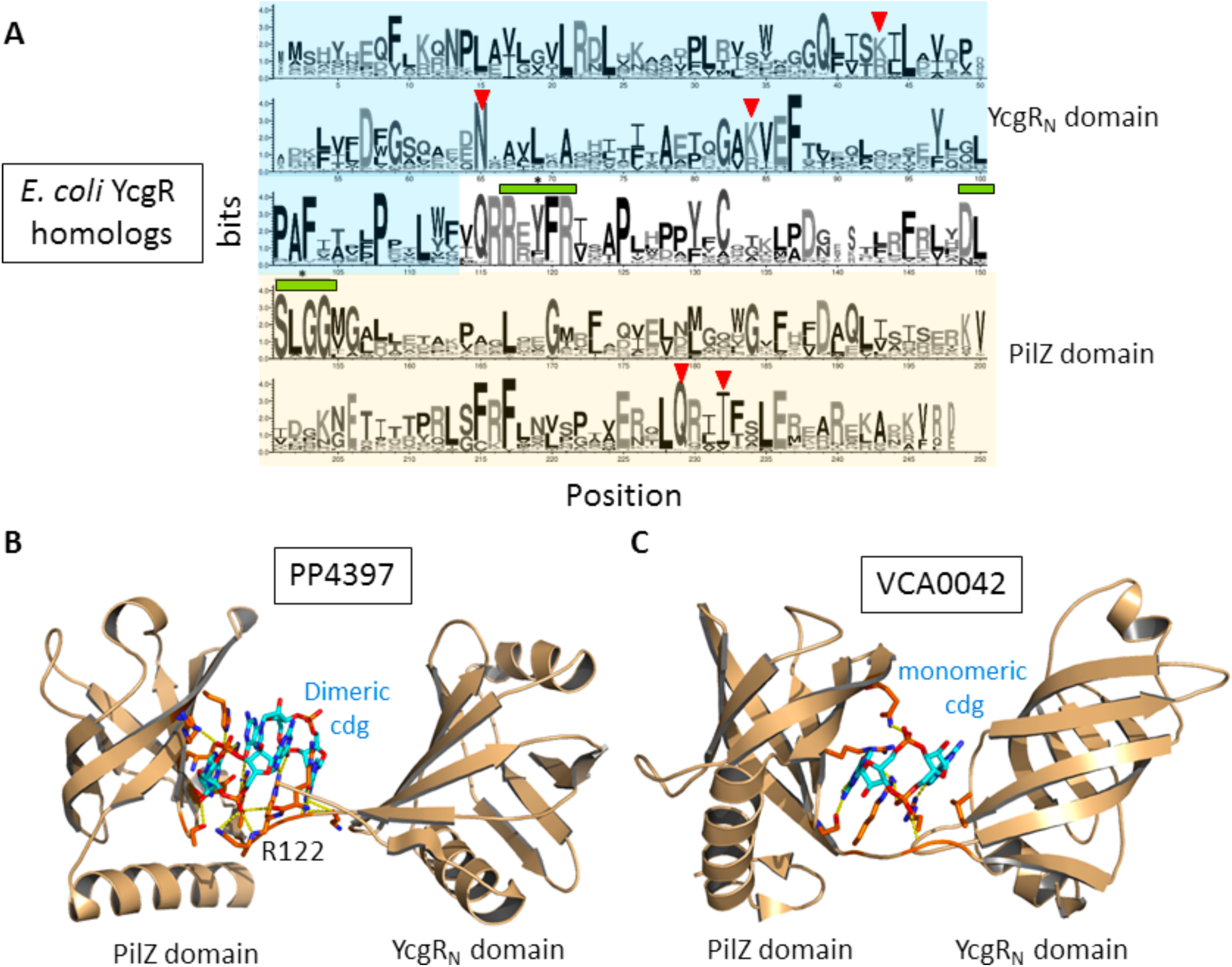
YcgR-type PilZ proteins. A. Sequence logo for the PilZ proteins in subcluster 1a, the subcluster that contains the prototype *E. coli* YcgR. The residues that are directly involved in binding FliG/FliM are indicated by the red triangles. B. Crystal structure of the YcgR-type protein PP4397 (PDB: 3KYF) with the residues involved in the direct binding of c-di-GMP labeled and shown as sticks. C-di-GMP is shown as sticks. C. Crystal structure of the YcgR-type protein VCA0042 (PDB: 2RDE) with the residues involved in the direct binding of c-di-GMP labeled and shown as sticks. C-di-GMP is also shown as sticks.

Cluster 4 contains 592 proteins that are mainly found in gamma-proteobacteria. This compact cluster contains the YcgR-type PilZ proteins FlgZ (also PA3353) from *P. aeruginosa* and PP4397 from *P. putida*. In contrast to YcgR, FlgZ was reported to interact with the stator protein MotC to influence swarming motility ^45–48^. Considering the different protein partners and cellular function for YcgR and FlgZ, they served as important references for generating isofunctional clusters. An alignment score of 25 allowed the segregation of YcgR and FlgZ into different clusters while retaining all the proteins with FliG/FliM-interacting residues within cluster 1. Cluster 4 also contains PP4397, a structurally characterized YcgR-like protein from *P. putida*. The crystal structure of PP4397 shows that di-domain proteins bind c-di-GMP similarly as single-domain PilZ proteins and reveals the β-barrel fold of the YcgR_N_ domain ^6^. The residues stemmed from the N- and C-terminal domains, as well as the c-di-GMP-binding motif (RxxxR) from the inter-domain linker (also termed as “c-di-GMP switch”), are mainly responsible for c-di-GMP binding ^4^ (Figure 10B). PP4397 binds two c-di-GMP molecules and mutational studies showed that the R122 residue of PP4397 is crucial for binding the dimeric c-di-GMP ^4^. Although the physiological role of PP4397 has not been established, the clustering of PP4397 with FlgZ indicates that PP4397 may bind to the MotC homolog in *P. putida* to regulate flagellum-dependent motility.

Several other clusters are also composed of YcgR-type proteins that likely to play different cellular roles than YcgR and PXO_00049. Cluster 5 is comprised of 562 YcgR-type proteins with the majority of them coming from the orders of *Bacillales* and *Lactobacillales.* This relatively compact cluster contains YpfA (also by the name DgrA), the sole YcgR-type protein in *B. subtilis*. YpfA was shown to suppress motility in a c-di-GMP dependent manner by interacting with the stator MotA ^49^. Cluster 6 consists of 520 PilZ sequences from a variety of genera and is one of the most “scattered” clusters. Cluster 7 comprises 221 YcgR-like PilZ proteins from 33 genera; whereas clusters 8 and 9 are composed of PilZ proteins that are mainly from the *Vibrio* genus. Cluster 8 contains the *V. cholerae* protein PlzD (VCA0042), the first di-domain PilZ protein to have the crystal structure determined ^4^. In contrast to PP4397 discussed above, PlzD binds monomeric c-di-GMP, rather than dimeric c-di-GMP, largely due to the replacement of an R/K residue located immediately upstream of the “RxxxR” motif (Figure 10C). The last cluster, cluster 9, contains the functionally unknown YcgR-type protein PlzE from *V. cholerae*.

Overall, the SSN analysis reveals that YcgR-type PilZ proteins are the most abundant di-domain PilZ proteins. Despite the fact they all contain a c-di-GMP-binding C-terminal PilZ domain and an N-terminal YcgR_N_ domain, divergent evolution has generated distinct subfamilies of YcgR-type proteins that are characterized by different binding partners (e.g. FliG, FliM, MotA and MotC etc.) and physiological functions. Such information will be instrumental in future studies of the YcgR-like proteins.

### Di-domain PilZ proteins that contain tandem PilZ domains

A few clusters of di-domain PilZ proteins (clusters 3, 14 and the MrkH cluster) contain an additional PilZ domain to form tandem PilZ proteins. Typically, one of the two PilZ domains is a canonical c-di-GMP-binding domain; whereas the other PilZ domain usually lacks the RxxxR and D/NxSxxG motifs and exhibit low sequence similarity.

A cluster that is predicted to have the PilZ-PilZ architecture is the MrkH cluster. This small cluster contains the *K. pneumonia* protein MrkH, which is the only tandem PilZ protein with known structure and cellular function. MrkH is a c-di-GMP-binding transcriptional regulator that regulates the expression of *mrk*ABCDF genes to control type 3 fimbriae and biofilm formation ^50^. The crystal structure of MrkH shows that the RxxxR and D/NxSxxG motifs are from the canonical C-terminal PilZ domain, and that the N-terminal PilZ domain also contributes to c-di-GMP binding through residues H69, G73 and R108 (Figure 11A). MrkH interacts with DNA mainly through a C-terminal helix in a c-di-GMP-dependent fashion^50^. Several positively charged residues from the C-terminal helix (R217, R218, K21, R232, K233, K234) and two residues (Y202, Q203) from elsewhere were involved in DNA binding. The DNA-binding residues of MrkH are conserved in all the proteins from this small cluster, indicating that the proteins are likely true functional homologs/orthologs of MrkH.

**Figure 11.**
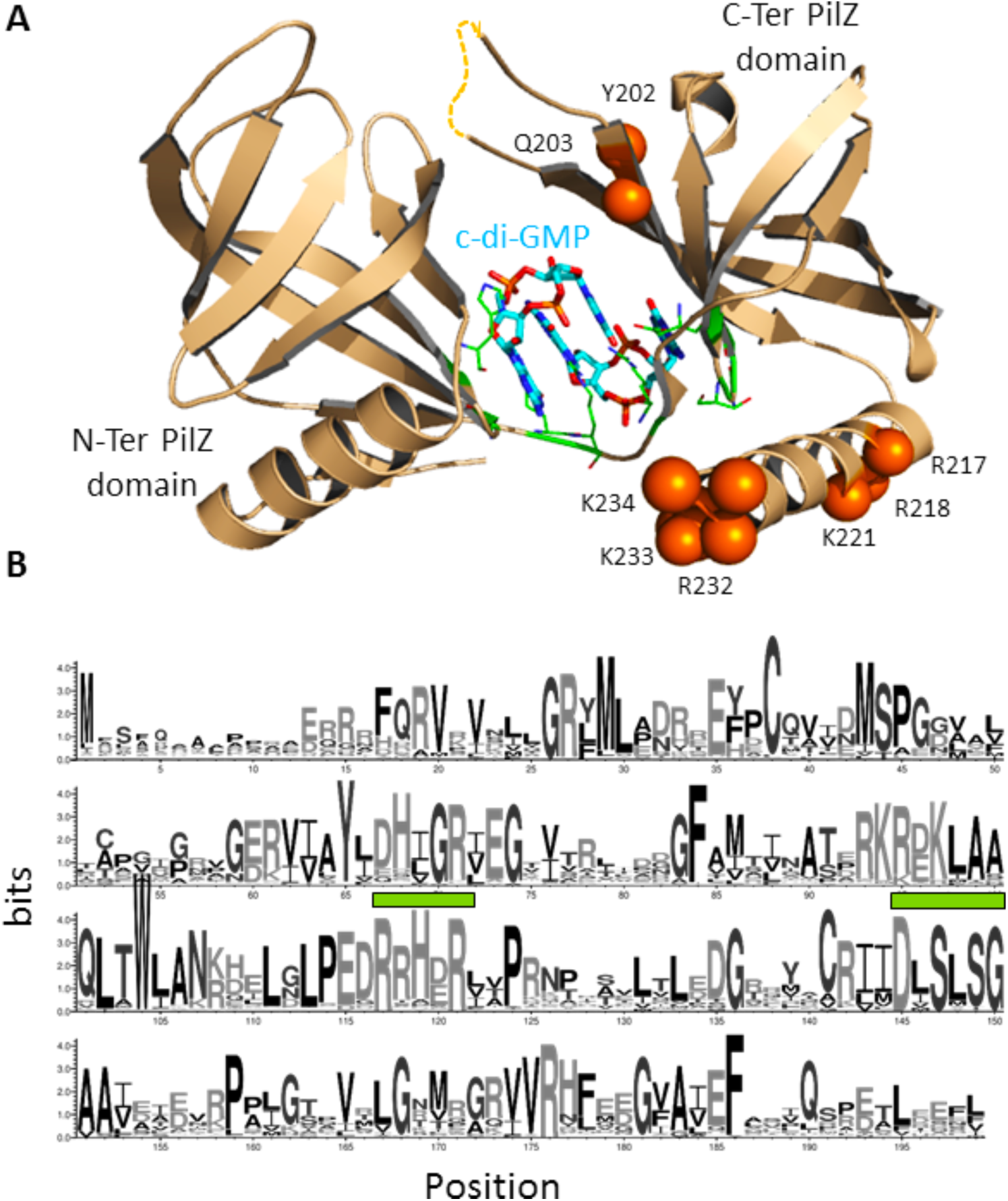
Di-domain PilZ proteins that feature a PilZ-PilZ architecture. A. Crystal structure of MrkH (PDB: 5KGO) with c-di-GMP-interacting motif colored in green and the α-carbon of the residues for DNA-binding shown as spheres. B. Sequence logo for the proteins from cluster 3, the largest cluster that contains PilZ-PilZ proteins. The two conserved c-di-GMP-binding motifs are indicated by the green bars. The absence of the DNA-binding residues observed in MrkH homologs indicates that the proteins are likely to have different cellular functions.

On the other hand, although proteins from clusters 3 and 14 account for 8% of the total di-domain PilZ proteins, none of the proteins has been characterized experimentally. The sequence logo for cluster 3 proteins shows that they contain the two c-di-GMP-binding motifs, but lack the DNA-binding residues of MrkH (Figure 11B) to indicate the non-canonical PilZ domain of the proteins may have a different function.

### Di-domain PilZ proteins with an inserted two-helix domain

The PilZ protein XCC6012 (also XCCB100-2234) from *Xcc* strain 17 is included in a small cluster (cluster 11) in the network. XCC6012 lacks the RxxxR and D/NxSxxG binding motifs and does not exhibit c-di-GMP binding ability ^23^. The crystal structure of XCC6012 shows that XCC6012 contains a central PilZ domain and two additional helices (α2 and α3, a total of 57 residues) inserted between the PilZ β1 and β2 strands^23^ (Figure 12A). XCC6012 exists in solution as a homotetramer stabilized by a central helical bundle formed through the helices α2 and α3 (Figure 12B) ^23^.

**Figure 12.**
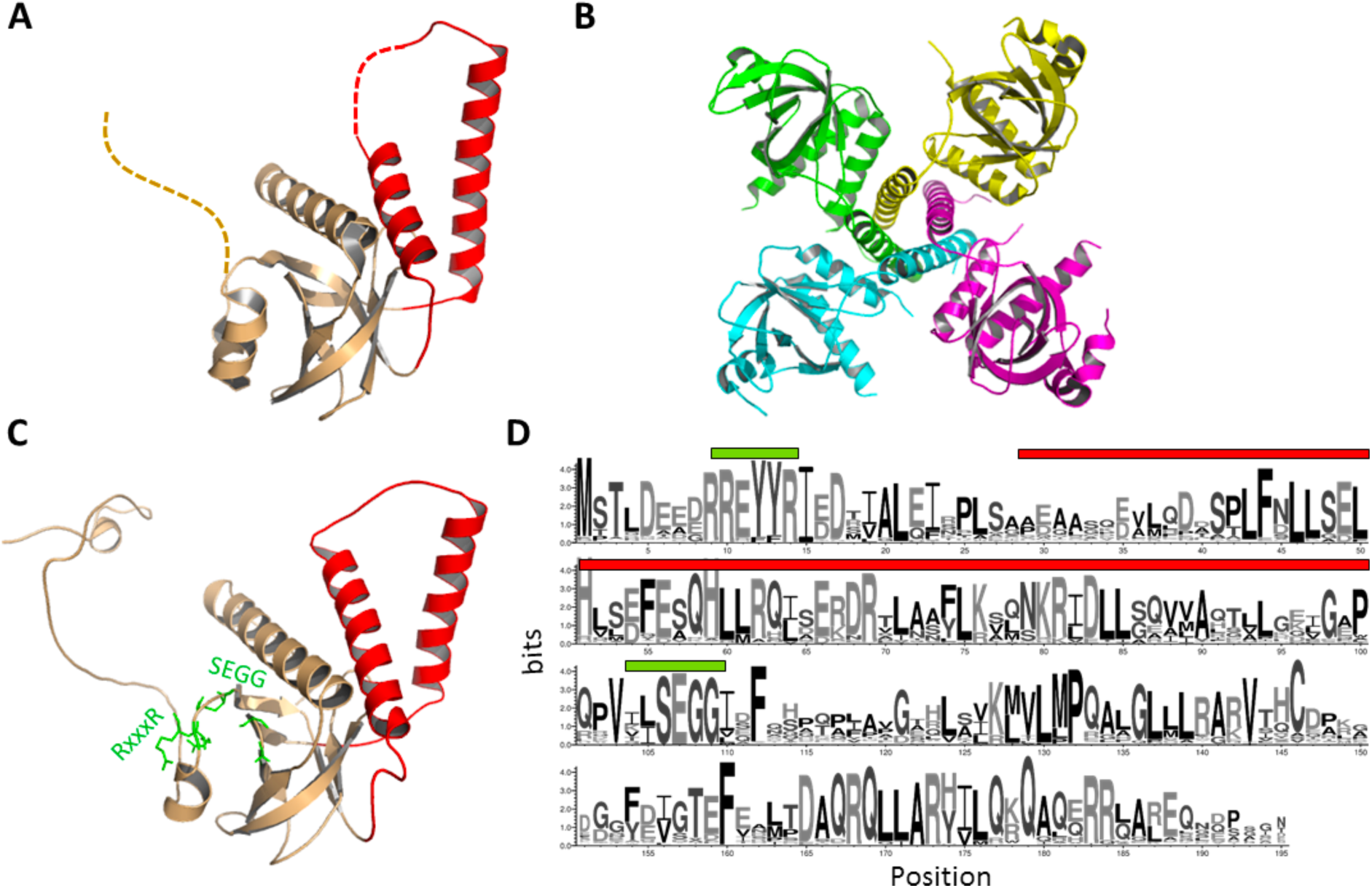
A unique family of PilZ proteins formed by domain insertion. A. The crystal structure of the subunit of XCC6012 shows the insertion of two long α-helices (In magenta) between the β1 and β2 strands of the PilZ domain. B. The crystal structure of the tetrameric XCC6012 shows the inserted helices is likely to drive the formation of tetramer. C. Structural model of PA2989 (a protein from cluster 2) shows a similar structure as XCC6012 with two inserted α-helices between the β1 and β2 strands. D. The sequence logo for the proteins of cluster 2 shows while the RxxxR motif is retained, the D/NxSxxG motif is replaced by the xxSEGG motif. The motifs are indicated by the green bar and the inserted helices are indicated by the magenta bar.

Proteins from two more clusters (clusters 2 and 12) are predicted to contain a similarly inserted helical domain. The second-largest cluster in the SSN (Cluster 2) is comprised of 659 proteins and contains a group of PilZ proteins with unknown function, including the *P. aeruginosa* protein PA2989. The cluster is divided into two sub-clusters, with the larger sub-cluster (2a) consisting of 618 proteins and the smaller subcluster 2b comprised of 43 proteins. Our structural modeling study on PA2989 suggests that this group of proteins also contain an inserted two-helix motif or domain between the PilZ β1 and β2 strands (Figure 12C). As a result of the sequence insertion, there is a large separation (>90 residues) between the RxxxR and SEGG motifs, in contrast to the ∼20-30 residue separation in other canonical PilZ proteins. In contrast to the proteins of cluster 11, the RxxxR motif for c-di-GMP binding can be found in this cluster of proteins. While the second motif D/NxSxxG can be found in the cluster 2b proteins, the motif is replaced by a slightly different xxSEGG motif in cluster 2a proteins. The replacement of (D/N)xSxxG by the xxSEGG motif does not seem to negatively affect c-di-GMP binding because PA2989 still can bind c-di-GMP with high affinity (K_D_ = 2.88 uM) ^51–, 52^. Meanwhile, the proteins from the small cluster 12 contain both RxxxR and (D/N)xSxxG motifs and are predicted to bind c-di-GMP. Together, the observations suggest that the proteins from clusters 2 and 11 originated from a domain insertion event and represents a distinct group of PilZ proteins with a slightly different c-di-GMP-binding motif.

### Other notable clusters of di-domain PilZ proteins

Apart from the clusters discussed above, the 163 proteins from cluster 10 contain an N-terminal domain recognized as a signal transduction response regulator domain. The scattered nodes may reflect a divergent evolution of the domains to interact with different histidine kinases. The compact cluster 15, which comprised of 97 proteins, is predicted to have a C-terminal HTH type DNA binding domain, indicating potential roles of the proteins in gene expression.

The PilZ protein PlzA (BB0733 from *B. burgdorferi*), which contains a structurally unknown N-terminal domain, does not cluster with any other proteins in the SSN. PlzA is a multifunctional protein that regulates bacterial motility and virulence by controlling gene expression in *B. burgdorferi* ^53–55^. The PilZ domain of PlzA contains the two c-di-GMP binding motifs, which is consistent with the observation that PlzA binds c-di-GMP with a *K*_D_ of 1.25 µM. Many non-YcgR di-domain proteins in other small clusters feature yet-to-be annotated C-terminal domains and will not be discussed further here.

## Discussion

As of today, only a small number of single- and di-domain PilZ proteins have been experimentally characterized. The scarcity of information about the cellular function and binding partner of PilZ proteins is in stark contrast to the impressive knowledge accumulated for non-PilZ c-di-GMP effectors ^56–57^. The vast number of PilZ proteins makes it pivotal to infer their function based on the sequence-structure-function relationship. As a step towards this goal, we explored the sequence and function space occupied by single- and di-domain PilZ domain proteins. To our knowledge, this is one of the first few studies that SSN analysis was applied to a non-enzymatic protein superfamily. Although the SSN tool was created originally to probe the scope of function diversification in enzyme superfamilies, the underlying assumption that sequence similarity generally correlates with protein function (e.g. catalytic mechanism, substrate specificity, ligand-binding and protein-binding) should remain valid for non-enzymatic protein superfamilies. Similar to SSN-based studies on enzyme superfamilies, our objective was to generate an overview of the function diversification to enable the formulation of reasonable hypotheses for experimental validation. For hypothesis generation, an SSN is most valuable when nodes are segregated into isofunctional clusters. The reference nodes or functionally characterized PilZ proteins were instrumental in guiding the segregation of nodes into putative isofunctional clusters. The formation of isofunctional clusters is also bolstered by the observation that functionally important residues are generally conserved in the putative isofunctional clusters or subclusters. Hence, we believe that the SSN analysis provides a rare glimpse into the scope of the sequence and function divergence of PilZ proteins.

One of the most notable realizations from the SSN analysis is that the single-domain PilZ proteins exhibit enormous sequence divergence despite the conserved β-barrel fold of the PilZ core. Variation of length and secondary structure in the C-terminal motif contributes to the sequence divergence in single-domain PilZ proteins. Considering the role of the C-terminal motif in binding protein partners as revealed by the studies on MapZ ^7^, the sequence variation in the C-terminus is likely to reflect a diversity of interacting protein partners. Evolving new C-terminal motifs to interact with different proteins, DNA/RNA partners to regulate different pathways could be one of the major forces driving the divergent evolution of PilZ proteins. The cellular function for the proteins from many of the clusters, including some large clusters, remains completely unknown today. For example, clusters 3 contains a large number of putative functional homologs with none of the proteins characterized. Given their wide occurrence, we propose that proteins from the large functionally unknown clusters be given the priority for future study.

An important revelation about single-domain PilZ proteins is that evolutionary divergence has generated PilZ proteins with different c-di-GMP binding affinity and binding mode. For the canonical single-domain PilZ proteins, the residues within or near the two c-di-GMP-binding motifs vary considerably (Figure 4). It was known that a change of the residue (R/K or L) located immediately upstream of the RxxxR motif can result in the binding of monomeric or dimeric c-di-GMP ^6^. Replacement of other residues in the vicinity of the two motifs could account for the large range of c-di-GMP binding affinity observed for the PilZ proteins^51^. More drastic divergence led to the loss of the c-di-GMP binding motifs to generate non-canonical PilZ proteins. Although the existence of non-canonical PilZ proteins is known, it is surprising that they constitute more than 17% of the single-domain proteins (clusters 1, 5, 6 and 18). As the largest of the non-canonical PilZ protein clusters that contains the homologs of PilZ (PA2960), cluster 1 may contain a group of proteins that regulate the assembly and function of Type IV pili. The proteins from clusters 5, 6 and 18, are likely to interact with different protein partners and thus have different biological functions, considering that the proteins possess different sets of conserved residues and lack the residues for FimX or PilB binding.

The most salient feature of the di-domain PilZ SSN is that the YcgR-type proteins far outnumber other di-domain PilZ proteins. In the SSN, about 48% of the proteins are YcgR-type proteins. The three experimentally characterized YcgR-like proteins YcgR (*E. coli),* FlgZ *(P. aeruginosa)* and YpfA *(B. subtilis)* have been reported to interact with FliG/FliM, MotC, and MotA respectively ^58^. Hence, it is apparent that the YcgR-type proteins have diverged to interact with different flagellar proteins to allow c-di-GMP to exert their control through distinct mechanisms. Sequence variation seems to have taken place in both the PilZ and YcgR_N_ domain. This is not surprising considering that both domains contribute to c-di-GMP-binding and both domains are involved in the interaction with the protein partners. As some of the YcgR-type proteins are not in the same cluster as the *E. coli* YcgR or *P. aeruginosa* FlgZ, they are likely to have different biological functions and it will be interesting to know whether they also regulate the flagellar motor activity by binding to other proteins than FliG/M and MotA/C.

Apart from the YcgR-type proteins, the SSN includes many clusters of poorly characterized di-domain PilZ proteins harboring a non-YcgR_N_ protein domain. Some of the non-YcgR_N_ domains can be annotated based on structural homology (e.g. REC domain, DNA binding domain), whereas many of the relatively small clusters feature unannotated N-terminal domain. The SSN includes a few clusters that contain two PilZ domains, featured by a canonical PilZ domain and a non-canonical PilZ domain. Grouping of the tandem PilZ proteins into different clusters indicates a functional divergence for some of the proteins from MrkH. Another interesting and unique group of di-domain proteins is this large cluster of proteins represented by the *P. aeruginosa* protein PA2989, which consists of a core PilZ domain and an inserted two-helical domain. The α-helical domain is likely to be a result of a gene insertion event in evolution.

In summary, we applied sequence similarity analysis to the PilZ protein superfamily to probe the scope of function divergence in the single- and di-domain PilZ proteins. The SSNs provide a global view of the sequence-function relationships and identify a large number of homologs/orthologs of structurally or functionally known PilZ proteins. For the proteins that do not cluster with functionally characterized PilZ proteins, we grouped them into putative isofunctional clusters to provide an overview of their taxonomical distribution and sequence features. As one step towards understanding the diverse roles played by the versatile PilZ domain in c-di-GMP-dependent signaling pathways, we hope the work will facilitate the annotation of the vast number of PilZ proteins encoded by bacterial genome and help to prioritize functionally unknown PilZ proteins for future studies.

## Materials and Methods

### Mining of PilZ protein sequences

PilZ protein sequences were collected from the InterPro database (v64.0) and UniProt Knowledgebase (Uniprot, July 2017 version). The sequences include the entire IPR009875 and IPR031800 families along with the FASTA sequences of a small number of PilZ proteins that are missing in the two families, such as PlzA^59^ (Uniprot ID: O51675) and MrkH^60^ homologs (Uniprot ID: G3FT00)), to yield a collection of 28,223 PilZ protein sequences. A histogram for the PilZ proteins according to the protein length was generated using the Enzyme Function Initiative and Enzyme Similarity Tool (EFI-EST) sever^15^ to group the proteins into single-domain (80-160 residues), di-domain (170-280 residues) and multi-domain proteins (>280 residues).

### Construction of sequence similarity networks (SSN) for single- and di-domain PilZ proteins

The SSNs were generated using EFI-EST sever (Option C), which accepts InterPro/Pfam numbers and user-defined FASTA sequence files as input (http://efi.igb.illinois.edu/efi-est/stepa.php) ^15^. From the 28,223 PilZ sequence collection, the protein sequences that contain 80-160 residues were submitted to the EFI-EST server to generate the SSNs for single-domain PilZ proteins; and the protein sequences that contain 170-280 residues were submitted to generate the SSNs for di-domain PilZ proteins. An alignment score corresponding to 35% and 40% sequence identity were used to generate the initial SSNs for single- and di-domain PilZ proteins respectively. Given the difficulty of visualizing the large number of nodes in the full network, representative node networks that contain “meta-nodes” generated by the EFI-EST sever were used for visualization and further analysis. In the representative node networks, sequences that share higher than 90% sequence identity are merged into one single “meta-node”. A more detailed description of how the SSNs are generated can be found in early publications ^15–16, 61^.

All networks were visualized in Cytoscape (v3.5.1) using the “organic” layout whereby nodes are clustered more compactly if they are more connected^62^, with more connection (or “edge”) within clusters indicate greater similarity. In this layout, the relative edge lengths were shown to be close to the BLAST E-value distances^16^. The alignment scores chosen for visualization of the PilZ SSNs were determined by stepping through alignment score values in annotated networks until major clusters containing known PilZs of distinct function (“reference nodes”) were separated from other unrelated clusters. The alignment score at which clusters were sufficiently distinct but not overly fractionated was chosen to avoid missing any potential relationships between the nodes.

The SSNs generated for single- and di-didomain PilZ proteins are included in the supplementary material as Cytoscape files. The reader can download the files and retrieve the sequences and other information associated with the protein name and sequence.

### Generation of protein sequence logos

Protein sequence logos were generated using the Weblogo tool^63^ (http://weblogo.threeplusone.com/) by inputting the multiple sequence alignment files that contain the protein sequences of individual SSN clusters (or sub-clusters). To retrieve the protein sequences, the cluster of interest was first created as a separate SSN in Cytoscape. The nodes within this new network were then exported into a table that contains the Uniprot ID of proteins and additional information represented by this network. The Uniprot IDs were used to retrieve the sequences from Uniprot and then used for Multiple Sequence Alignment using Clustal Omega with default settings (http://www.ebi.ac.uk/Tools/msa/clustalo/)^64^. Gaps in the sequences logs were removed manually. All the sequence logo files will be available to the readers upon request.

### Homology modeling

The structural model for the di-domain PilZ protein PA2989 was constructed by homology modeling using the crystal structure of XCC6012 (PDB: 3PH1) as the template. The homology model was generated using I-TASSER (Iterative Threading Assembly Refinement, https://zhanglab.ccmb.med.umich.edu/I-TASSER/^65^), which constructs full-length atomic models by homology threading and iterative template fragment assembly simulations. The crystal structure of XCC6012 (PDB: 3PH1) was identified by an automatic template search and used as the template for homology modeling. The C-score of the PA2989 structural model shown in this manuscript is −2.57. An estimated TM-score of 0.42±0.14 and RMSD of 11.8±4.5Å were found between the template and the structural model.

## Author contribution

Z.X.L., Q.W. C., and L. X. conceived and initiated the study. Q. W. C. and S. S. performed data mining structural modeling and bioinformatic analysis. Q. W. C and Z.X.L. wrote the manuscript.

## Acknowledgment

This work is supported by an SBP-01 grant (Z.-X. L.) from NRF, Singapore. This work was also supported by the Introduction of Innovative R&D Team Program of Guangdong Province (X. Xu., No. 2013S034).

